# Inferring cellular trajectories from scRNA-seq using Pseudocell Tracer

**DOI:** 10.1101/2020.06.26.173179

**Authors:** Derek Reiman, Heping Xu, Andrew Sonin, Dianyu Chen, Harinder Singh, Aly A. Khan

**Author notes:** These authors contributed equally (Derek Reiman, Heping Xu).

## Abstract

Single cell RNA sequencing (scRNA-seq) can be used to infer a temporal ordering of dynamic cellular states. Current methods for the inference of cellular trajectories rely on unbiased dimensionality reduction techniques. However, such biologically agnostic ordering can prove difficult for modeling complex developmental or differentiation processes. The cellular heterogeneity of dynamic biological compartments can result in sparse sampling of key intermediate cell states. This scenario is especially pronounced in dynamic immune responses of innate and adaptive immune cells. To overcome these limitations, we develop a supervised machine learning framework, called Pseudocell Tracer, which infers trajectories in pseudospace rather than in pseudotime. The method uses a supervised encoder, trained with adjacent biological information, to project scRNA-seq data into a low-dimensional cellular state space. Then a generative adversarial network (GAN) is used to simulate pesudocells at regular intervals along a virtual cell-state axis. We demonstrate the utility of Pseudocell Tracer by modeling B cells undergoing immunoglobulin class switch recombination (CSR) during a prototypic antigen-induced antibody response. Our results reveal an ordering of key transcription factors regulating CSR, including the concomitant induction of *Nfkb1* and *Stat6* prior to the upregulation of *Bach2* expression. Furthermore, the expression dynamics of genes encoding cytokine receptors point to the existence of a regulatory mechanism that reinforces IL-4 signaling to direct CSR to the IgG1 isotype.

## INTRODUCTION

Single-cell RNA sequencing (scRNA-seq) has emerged as a dominant tool for analyzing the transcriptional states of individual cells in diverse biological contexts (Regev et al., 2017; Schaum et al., 2018). Computational analyses of scRNA-seq datasets have enabled rigorous delineation of known cellular identities as well as the discovery of novel cell types (Neu et al., 2017). Such datasets have also been used to infer a temporal ordering of dynamic cellular states or cellular trajectories (Saelens et al., 2019). For example, the field of immunology has benefited significantly from the adoption of scRNA-seq in order to characterize cellular states in the context of development and differentiation of distinct innate and adaptive lineages (Kakaradov et al., 2017; Olsson et al., 2016; Yu et al., 2016), including responses to various perturbations (Bossel et al., 2019; Dixit et al., 2016; Neu et al., 2019), as well as in immune system diseases (Lönnberg et al., 2017; Zhang et al., 2019; Zheng et al., 2017). In spite of the tremendous progress, the inference of cellular trajectories from scRNA-seq datasets remains challenging when analyzing heterogeneous cellular compartments with complex dynamics.

Current computational methods for cellular trajectory inference rely on two crucial steps. In the first step, dimensionality reduction techniques (Sun et al., 2019), such as PCA (Street et al., 2018), ICA (Trapnell et al., 2014), and UMAP (Wolf et al., 2019), are used to project and visualize single cells based on their gene expression profiles in low dimensional space (Figure 1A, left). While single cell transcriptional profiles have high dimensionality due to the thousands of genes profiled, their intrinsic dimensionalities are typically much lower. Gene expression during a biological process can be directed by small combinations of transcription factors that regulate large gene modules in a time-dependent manner. Thus, unsupervised low dimensional projections can reveal salient temporal structure in large-scale scRNA-seq datasets, especially when a dominant transcriptional regulatory program directs the biological process. In the second step of trajectory inference, pathfinding algorithms, such as minimum spanning trees (Street et al., 2018; Trapnell et al., 2014) or k-nearest neighbor graphs (Herring et al., 2018; Wolf et al., 2019), are utilized for inferring an ordering of cells in the low dimensional space (Figure 1A, center). The cells ordered in the inferred trajectory are typically mapped to a virtual temporal axis called “pseudotime”, which is bounded by two cells representing the start and end of the cellular trajectory. Gene expression abundances from the original high dimensional profiles can then be plotted along a pseudotime coordinate to display their changes along the inferred trajectory (Figure 1A, right).

**Figure 1:**
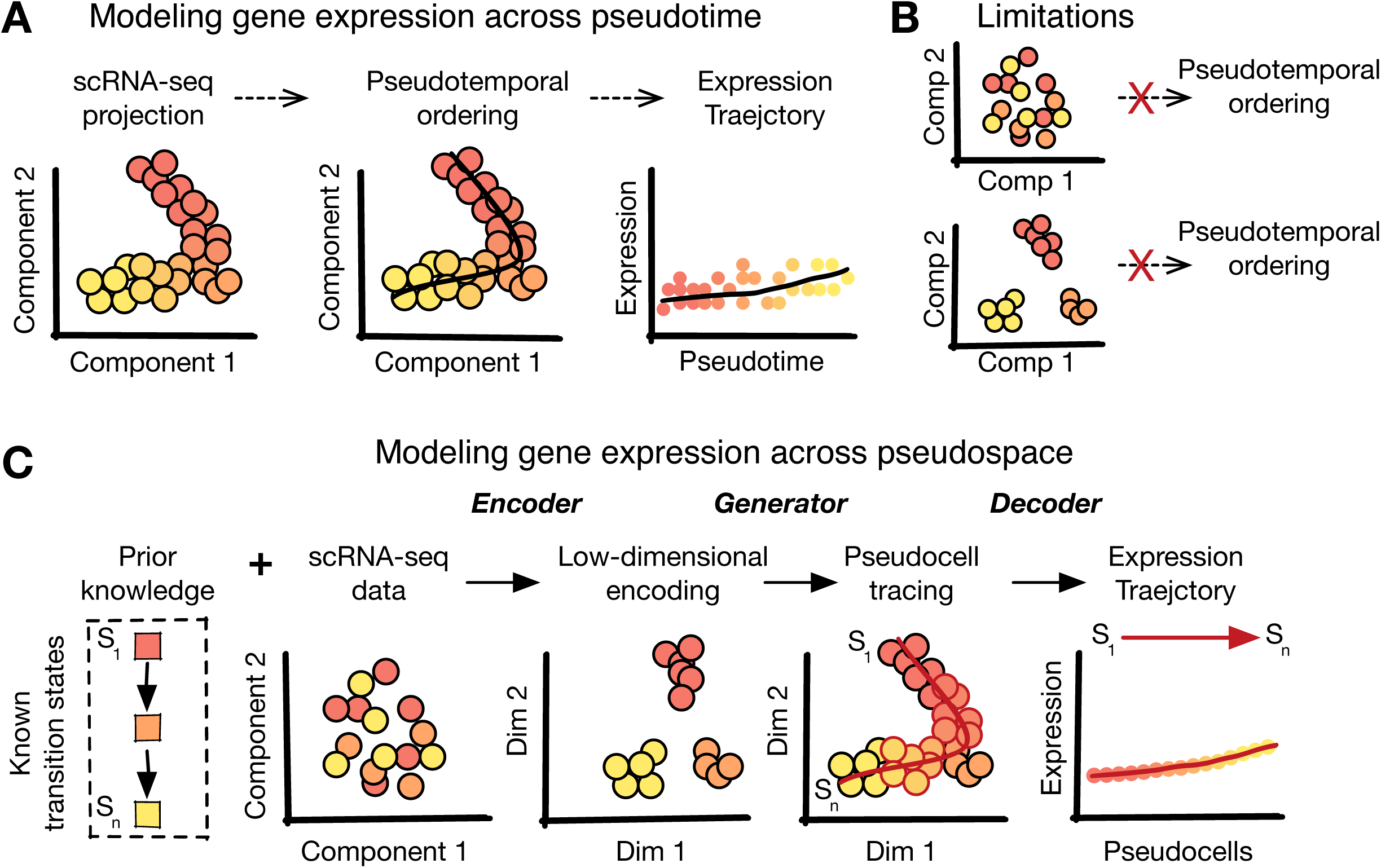
Pseudocell Tracer, a framework for modeling cellular trajectories in complex systems. (A) An overview of pseudotime trajectory inference. (B) Some scenarios that may obstruct pseudotime ordering. (C) Pseudocell Tracer. Given some prior knowledge about a model system, we aim to predict expression trajectories by generating pesudocells at regular intervals along a virtual cell-state axis, even though such cells may be sparsely captured in single cell profiling data.

Heterogeneous cellular compartments with complex temporal dynamics can present unique hurdles to trajectory inference. We consider two scenarios that are common within such complex systems and that limit the use of standard inference methods. In the first scenario, cells utilizing concurrent transcriptional regulatory modules, such as those controlling cell cycle, metabolism and differentiation, may not reveal the developmental trajectory of interest along any particular axis using unsupervised dimensionality reduction techniques (Figure 1B, top). Dimensionality reduction works by minimizing (or maximizing) some global statistical measure on the gene expression profiles, such as percent variance explained in each orthogonal dimension of PCA. Thus, there is no guarantee that any single unsupervised dimensionality reduction technique can uncover a specific temporal pattern of interest. In the second scenario, all transitioning cellular states along a given trajectory may not be equally populated, resulting in greater capture of some cell states and sparse capture of other states (Figure 1B, bottom). This non-uniform sampling occurs when cells do not follow a constant rate of development or differentiation during a time-dependent process. Consequently, this can result in the lack of an observable continuum of cell states or temporal structure in low dimensional space, hindering cell ordering. Thus, these scenarios illustrate some of the key hurdles to trajectory inference for complex cellular compartments.

Existing computational tools for analyzing scRNA-seq datasets typically do so without reference to any of the underlying biological guideposts of the system used to generate the data. We hypothesized that by developing algorithms that take advantage of validated prior biological knowledge, we could extract otherwise unresolvable trajectories from scRNA-seq datasets, especially when analyzing heterogeneous cellular compartments with complex dynamics. To test this hypothesis, we develop a supervised machine learning framework, called Pseudocell Tracer, that enables modeling of cellular trajectories in complex dynamical systems. We accomplish supervision by harnessing adjacent information about the underlying biological process. In most biological systems, there is some prior validated knowledge about the underlying cellular states, directionality and dynamics of the process, which can be integrated into a computational model. For example, cellular differentiation is often tracked by the level of expression of one or more specific markers such as cell surface proteins or regulators i.e., transcription factors. The expression level of such markers or regulators can reflect a “developmental clock” and therefore serve as an estimate of the progression between the progenitor and terminal differentiation state in a complex biological process. We refer to this kind of prior knowledge as adjacent biological information.

To implement Pseudocell Tracer we harnessed recent advances in deep generative modeling. Generative adversarial networks (GANs) can learn a latent space from which to simulate gene expression profiles of cells that are indistinguishable from a distribution of real cells (Marouf et al., 2020). In particular, it has been previously proposed that the interpolation of cells in the latent space may be a means for simulating pseudocells along some cellular trajectory (Ghahramani et al., 2018). Notably, GANs cannot directly shape the latent space, for example, to reflect prior knowledge about a complex cellular compartment. However, autoencoders can learn latent spaces for scRNA-seq that satisfy specific biological constraints (Eraslan et al., 2019). The integration of such models along with the use of adjacent biological information to supervise their training has not received significant attention. Pseudocell Tracer integrates such models and generates pseudocells along specific cellular trajectories in a stepwise process. First, Pseudocell Tracer uses an encoder (Figure 1C, left) to project a low dimensional representation of the scRNA-seq data, while remaining faithful to adjacent biological information. Then, the framework uses a generator (Figure 1C, center) to simulate pesudocells at regular intervals in the latent space by using the same adjacent biological information as a guide. Finally, these pseudocells are subjected to a decoder (Figure 1C, right) to observe gene expression dynamics along the trajectory and provide novel insights into the underlying regulatory mechanisms. Taken together, Pseudocell Tracer infers trajectories in “pseudospace” rather than in “pseudotime”.

We apply Pseudocell Tracer to the process of somatic DNA recombination that B cells of the immune system undergo upon antigen encounter. The process termed class switch recombination (CSR) results in exchange of the constant region (M isotype) of the immunoglobulin heavy chain protein (IgH) to one of several other isotypes thereby generating IgG, IgA and IgE antibody expressing cells (Manis et al., 2002; Stavnezer et al., 2008). B cells that switch their IgM locus to one of the other Ig isotypes, via DNA recombination, can be viewed as moving down distinct cellular trajectories since different cytokine signals and transcription factors have been shown to promote specific types of isotype switching. An understanding of the timing and expression of the various signaling components and transcription factors associated with distinct CSR trajectories remains to be thoroughly explored and has not been analyzed in vivo by single-cell transcriptional profiling. Using an scRNA-seq dataset generated in the context of a prototypic antigen-specific B cell response we demonstrate that standard trajectory inference methods fail to assemble appropriate CSR trajectories. Instead, Pseudocell Tracer trained with adjacent information in the form of relative expression of isotype-specific transcripts enhances both dimensionality reduction and trajectory inference. In so doing it reveals the relative timing and orchestration of key cytokine receptors and transcription factors regulating a particular CSR trajectory.

## RESULTS

### Experimental system and the scRNA-seq dataset

Mouse B cell responses to the model antigen 4-hydroxy-3-nitrophenylacetyl-keyhole limpet hemocyanin (NP-KLH) have been used to reveal fundamental principles underlying antibody isotype switching and affinity maturation (Furukawa et al., 1999; Jacob et al., 1991). To analyze the dynamic transcriptional states of activated B cells we performed full-length scRNA-seq on NP-specific germinal center B cells at the peak of the response (day 13) (Figure 2A). We visualized the scRNA-seq data using t-distributed stochastic neighbor embedding (t-SNE) (Figure 2B) and observed that it failed to distinguish cells on the basis of their isotype identities (Figure 2C).

**Figure 2:**
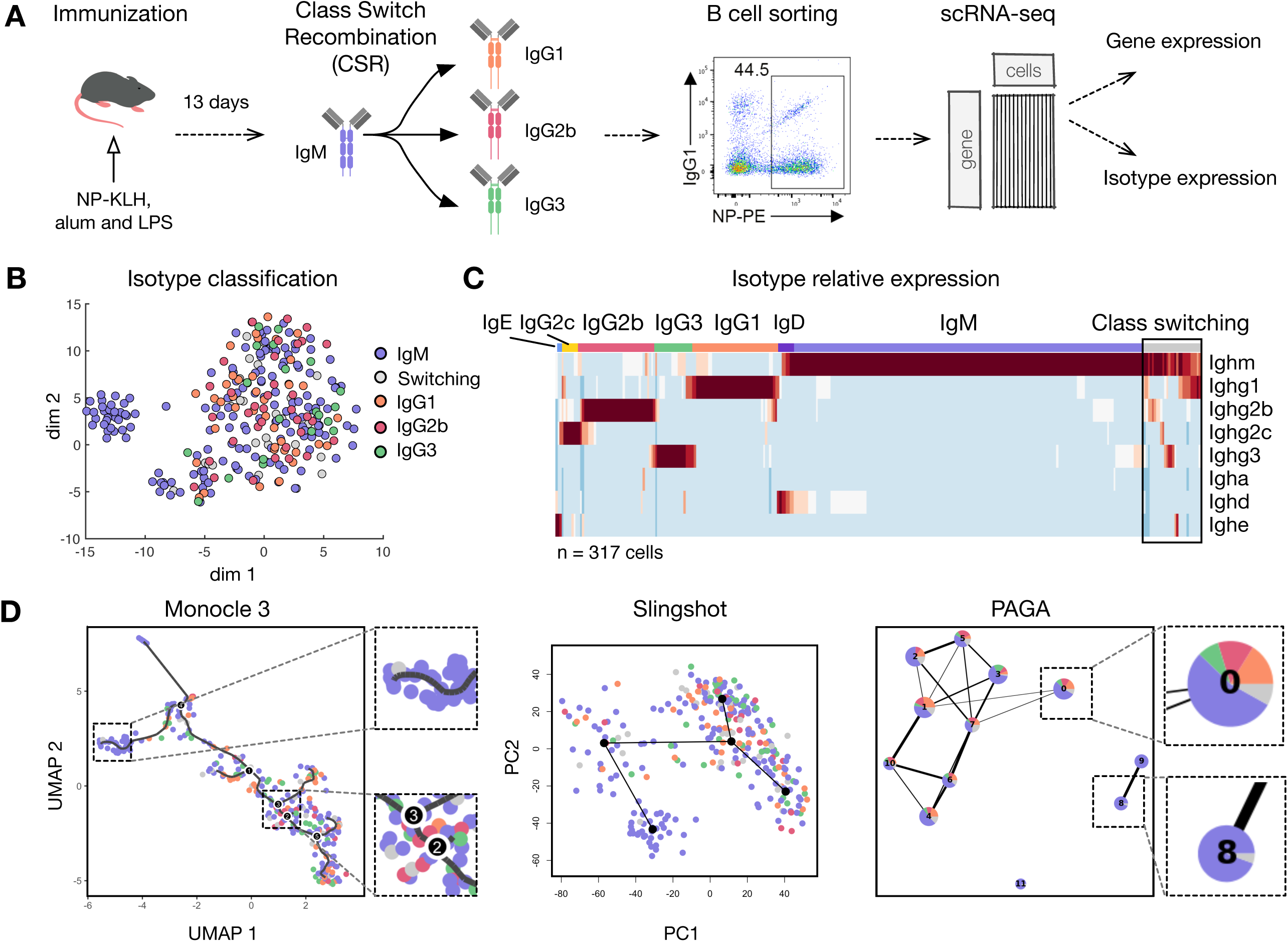
Class switch recombination process. (A) Overview of experimental system. (B) t-SNE of RNA-seq data, colored by isotype. (C) Relative isotype expression for all cells. N = 317. All isotype expressions sum to one for a given cell. C. Proportion of istoypes classified among all cells. (D) Output from Moncole3, Slingshot, and PAGA.

We reasoned that in B cells undergoing CSR, Ig transcripts encoding the M isotype would diminish whereas those encoding the switched isotype would increase. We calculated the relative isotype expression within a B cell by dividing the log2(TPM + 1) expression of each distinct isotype transcript (Ighm, Ighg1, Ighg2b, Ighg2c, Ighg3, Igha, Ighd, and Ighe) with the cumulative expression of all isotypes. Hierarchical clustering of relative isotype expression revealed 6 clusters of B cells, of which 5 clusters were dominated by a single isotype and reflected cells that had undergone CSR (Figure 2C). As expected for the immunization conditions noted above, IgM expressing B cells primarily switched their isotypes to IgG1, IgG2b, or IgG3 (Stavnezer et al., 2008). Notably, the clustering based on relative isotype expression values revealed a group of transitioning cells that were undergoing CSR from IgM to one of the following isotypes IgG1, IgG2b, IgG2c, IgG3 and IgE. As the relative isotype expression is expected to monotonically track the progression of CSR we reasoned that it would represent suitable information embedded within the dataset to enable deconvolution of the distinct CSR cellular trajectories within the transitioning B cells.

We therefore evaluated the utility of existing computational pipelines, in particular, Moncole3 (Qiu et al., 2017), Paga (Wolf et al., 2019), and Slingshot (Street et al., 2018) for delineating CSR trajectories within our dataset. These methods utilize a variety of dimensionality reduction techniques and temporal ordering methods. However, none of the methods recovered a coherent CSR trajectory that delineated a path from, for example, IgM to IgG1 (Figure 2D). In the case of Monocle3, the low dimensional projection of single cell data by UMAP was neither able to distinguish stably switched cells by their isotype nor cluster cells presumptively undergoing CSR. Similarly, the Slingshot and PAGA representations failed to distinguish cells on the basis of their CSR trajectories.

We hypothesized that the failure of these unsupervised dimensionality reduction techniques to uncover the CSR trajectories was due to other dominant dynamic gene expression programs in germinal center B cells, particularly involving the cell cycle. To evaluate this hypothesis, we analyzed the expression patterns of cell cycle regulators, which revealed clustering of cells based on their cell cycle phase (Supplementary Figure 1). Taken together, while our directed analysis of the scRNA-seq dataset revealed a minor cluster of transitioning B cells that were undergoing CSR to different isotypes, existing unsupervised methods were unable to reveal such cells as a distinct cluster(s) and therefore temporally order their distinct trajectories.

### Overview of Pseudocell Tracer

The core of the Pseudocell Tracer framework (Figure 3A) is based on the following two components that are used successively: (1) a supervised autoencoder to perform dimensionality reduction and (2) a conditional generative adversarial network (CGAN) to generate hypothetical cell states or pseudocells. The main difference between an unsupervised and supervised autoencoder is the additional information provided to facilitate learning a low dimensional projection (Figure 3A, left). Both unsupervised and supervised autoencoders function to encode high-dimensional data into a low-dimensional latent space. However, the supervised autoencoder aims to specifically learn an encoder that transforms the scRNA-seq data into a latent space that conforms to the adjacent information, relating to a specific biological context or process. In the context of modeling CSR, the latent space is shaped by relative expression of the different Ig constant region transcripts. Thus, individual B cells with similar relative isotype expression profiles will have similar latent encodings. The architecture of the supervised autoencoder contains both an encoder and decoder (Supplementary Figure 2). The encoder functions to project high-dimensional data into a low-dimensional latent space, which is shaped by the adjacent biological information (Figure 3B, top), and the decoder functions to reverse the low dimensional encoding of cells from the latent space (Figure 3B, bottom). When combined together, the encoder performs dimensionality reduction, while the decoder generates from the latent space a reconstruction as close as possible to the observed input data.

**Figure 3:**
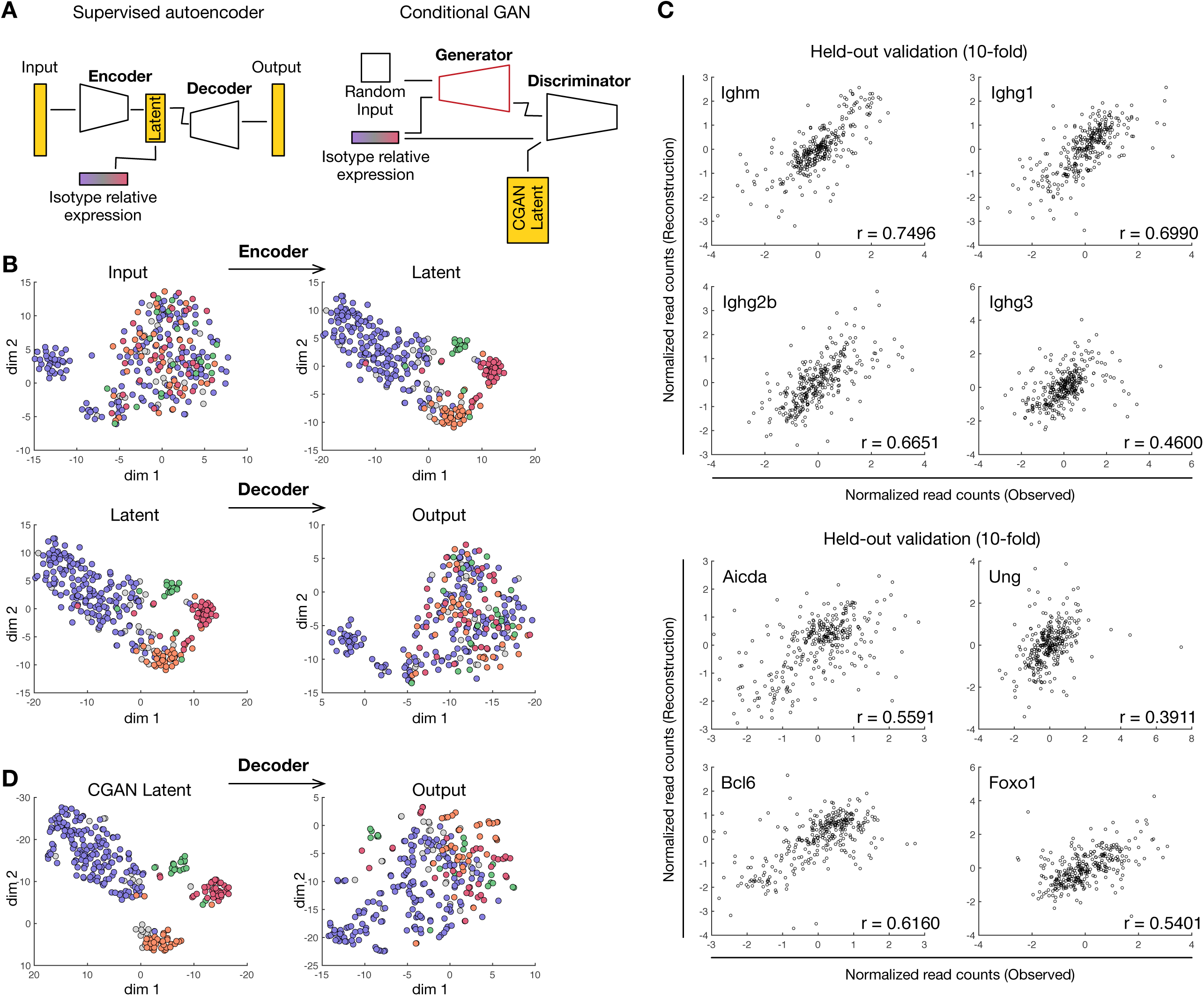
Pseudocell Tracer efficiently integrates adjacent biological information and accurately simulates gene expression profiles in pseudocells. (A) Overview of neural network model combining a supervised autoencoder with a conditional GAN. (B) t-SNE visualization of the input and output used in the supervised autoencoder; encoder (top) and decoder (bottom). (C) Scatter plot between observed and predicted expression values on held out cells. r denotes Pearson correlation between ground truth and predicted values. Isotype expression (top) and example CSR genes (bottom). (D) t-SNE visualization applied to cGAN prediction and subsequent output from decoder.

Visualization of the latent space for the scRNA-seq data revealed specific clustering of cells by their dominant isotype (Figure 3B, top). To characterize the robustness of the model on new or held out data, we evaluated the supervised autoencoder using 10-fold cross-validation (Figure 3C). For each partition, 90% of the data was used for training and 10% was set aside as a blind test. For training, an additional 10% of the training set was used for early stopping (see Methods). Once training finished, the test set was then encoded and decoded. Scatterplots of the held-out test predictions were generated for IgM and IgG isotypes, demonstrating high correlation between observed gene expression and the reconstructed expression. Similarly, other regulatory factors associated with CSR, demonstrated high correlation. Although inverse transformation of PCA can similarly approximate input data, the integration of adjacent biological information lacks a formal basis in PCA. Taken together, the supervised autoencoder successfully both learned an encoder for dimensionality reduction informed by relative isotype expression and a decoder for mapping low dimensional encodings back to full transcriptional expression profiles.

In the second step of our approach we trained a CGAN to simulate pseudocells (Supplementary Figure 3). Notably, the inference procedure for the generative model is performed in the latent space that is learned by the autoencoder. The use of the low-dimensional latent space is necessitated by the instability of GANs in high dimensional settings. Importantly, any CGAN simulation in low dimensional space can be easily mapped to the input space using the decoder and generate high dimensional full transcriptional profiles for individual cells. The main difference between a GAN and a CGAN is the conditional information associated with the generator and discriminator (Figure 3A, right). Both consist of two neural networks competing against each other such that one network, called the generator, seeks to produce realistic output data from a random input vector, and the other network, called the discriminator, is tasked with discriminating between the real and generated data. Importantly, the CGAN conditions the inference of both the generator and the discriminator on adjacent information. In the context of modeling CSR, the generator aims to simulate realistic latent encodings of cells that are conditioned on relative isotype expression profiles, in the same manner as the autoencoder in the first step. Thus, after fully training the CGAN, latent encodings for pesudocells can be simulated using the generator based on their relative isotype expression profiles. These latent encodings can then be subjected to the decoder utilized in the previous step to generate high dimensional transcriptional profiles of hypothetical cells that conform to the input data as well as the adjacent information that represents a key biological prior(s).

To qualitatively evaluate the representation learned by the generator, relative isotype expression and corresponding low-dimensional encodings from the scRNA-seq data were used to train the CGAN. The CGAN model was trained to equilibrium. Notably, only the discriminator directly observes low-dimensional encodings of the expression data while the generator improves its simulations through interaction with the discriminator. Pseudocells were generated for the observed relative isotype expressions, subsequently decoded, and visualized (Figure 3D). Visualization of the CGAN latent space reproduced specific clustering of cells by their dominant isotype (Figure 3B, top) and subsequent decoding reconstructed a complex and heterogeneous B cell landscape qualitatively similar to the real scRNA-seq data. Thus through sequential application of an autoencoder and a CGAN both conditioned with prior biological information, *Pseudocell Tracer* provided a supervised framework for generation of hypothetical B cells undergoing CSR.

### Pseudocell Tracing the CSR Process

We define a B cell IgH isotype trajectory based on a cellular progression from the IgM to an alternate IgH isotype. To demonstrate the utility of *Pseudocell Tracer* in inferring cellular trajectories that can be overwhelmed in complex and heterogeneous cellular compartments, we modeled the IgM to IgG1 class switch recombination process. First, we simulated a relative isotype expression profile with IgM at 100% and all other isotypes at 0%. For each cell-state increment along the IgG1 trajectory, we reduced the relative abundance of IgM by 1% and increased the relative abundance of IgG1 by 1%. We continued generating relative isotype expression profiles until IgG1 reached 100% and IgM reached 0% (Figure 4A). Overall, we simulated 101 points along the IgG1 trajectory. We then generated 100 latent encodings for each point using the previously trained CGAN in order to estimate a 95% confidence interval. Finally, we used the previously trained decoder to convert each latent encoding to a full transcriptional expression profile, resulting in 10,100 pseudocells which traced the progression from IgM to the IgG1 state within the trajectory.

**Figure 4:**
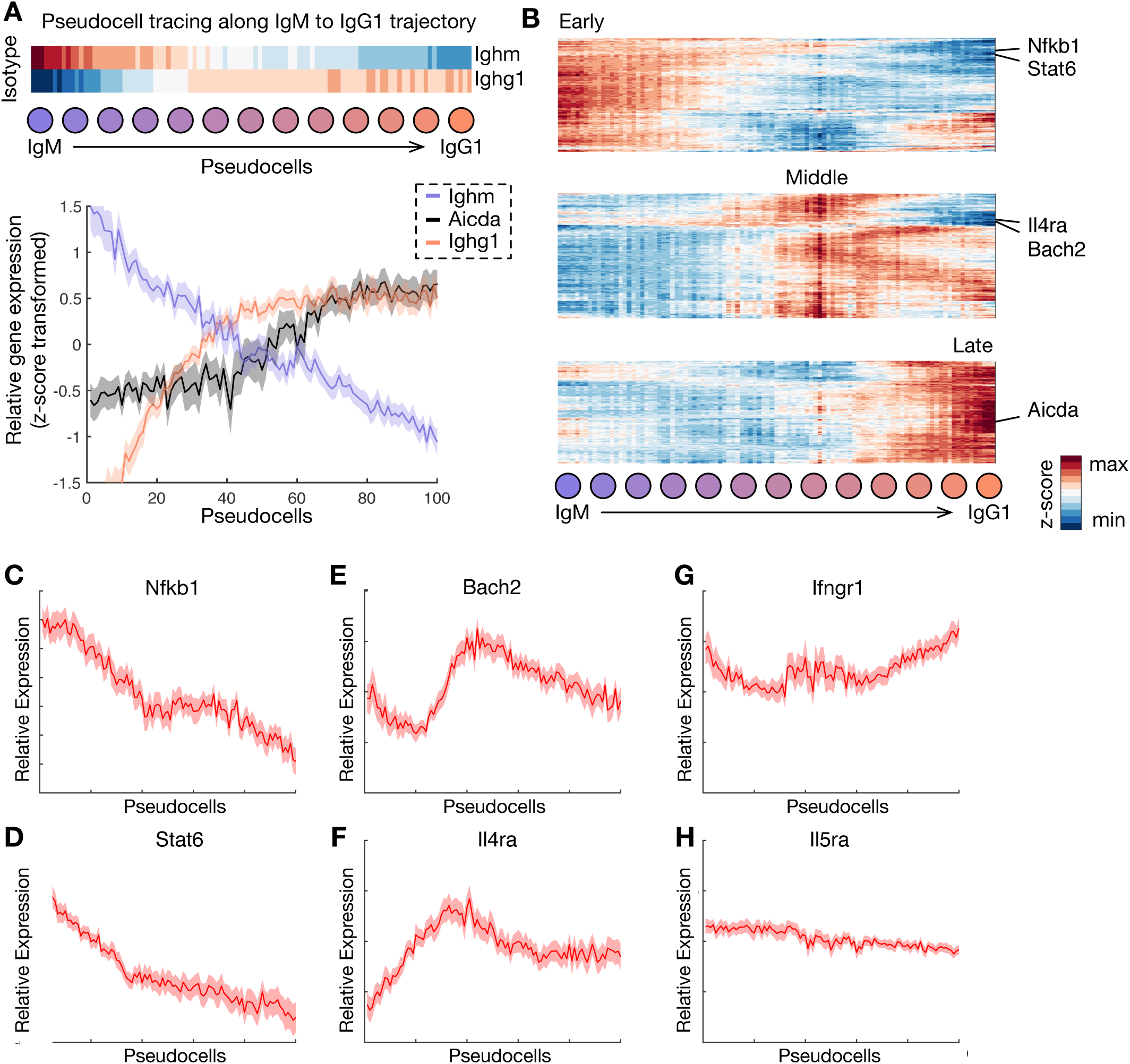
Pseudocell Tracer models IgG1 class switching process. (A) Pseudocells generated along the IgM to IgG1 axis. Heatmap of predicted *Ighm* and *Ighg1* gene expression changes (top), where each time point is an average of 100 simulations. Plot of relative expression of *Aicda, Ighm* and *Ighg1* along the IgM to IgG1 axis (bottom), where solid line indicates average expression and shading indicates 95% confidence interval. (B) Hierarchical clustering and segmentation of gene associated with CSR. Heamap of early (top), middle (center), and late (bottom) transcriptional dynamics are depicted. Plots of relative expression for key genes with specific dynamics, including (C) Nfkb1, (D) Stat6, (E) Bach2, and (F) Il4ra. Plots of relative expression for genes reflecting low variability throughout class switching, including (G) Ifngr1 and (H) Il5ra.

To determine if the pseudocell tracing of the IgG1 trajectory was consistent with known experimental findings, we examined the transcriptional dynamics of *Aicda* gene expression in relation to Ighm and Ighg1 transcripts. *Aicda* encodes the activation induced cytidine deaminase (AID) which is a direct mediator of the intrachromosomal IgH recombination events in B cells that result in CSR. We plotted the various relative expression profiles (z-score transformed) across the IgM to IgG1 pseudocell tracing (Figure 4A). As expected, we observed decreased relative expression of Ighm and increased expression of Ighg1 transcripts as pseudocells progressed from an IgM cell state to an IgG1 cell state. The *Aicda* transcript profile revealed a biphasic pattern. Despite a smaller number of IgG2 and IgG3 cells captured, similar relative expression profiles for *Ighg2, Ighg3* and *Aicda* were observed for other IgG trajectories (Supplementary Figure 4). The inferred biphasic expression profile of *Aicda* suggests a model where its initial levels in antigen-induced GC B cells are likely sufficient for promoting CSR. The increased levels at later timepoints in the CSR trajectory may function in promoting somatic hypermutation (SHM), another key molecular process required for affinity maturation that is directly mediated by AID (see Discussion).

To explore the regulatory underpinnings of IgG1 isotype switching, we assembled a comprehensive view of the gene expression dynamics across the IgG1 trajectory. We focused our analysis on dynamically expressed genes by selecting the top 2,000 most variable genes. Heatmap visualization of the relative expression profiles across the IgM to IgG1 pseudocell tracing revealed 3 granular transcriptional phases associated with this CSR trajectory, designated early, middle, and late (Figure 4B).

We characterized the dynamics of several key transcription factors that are implicated in regulating CSR within the phases. Genes induced within the early transcriptional phase included *Nfkb1* and *Stat6* (Figure 4C,D). *Nfkb1* knockout mice have lower serum IgG1 and IgE antibodies (William et al., 1995). Stat6 induces *Ighg1* germline transcription, an event that is obligatory for the intrachromosomal DNA recombination that leads to IgG1 switching (Harris et al., 1999). Furthermore, we observed an increase in *Bach2* expression during the middle transcriptional phase (Figure 4E). *Bach2* is required for both CSR and SHM and regulates *Aicda* expression (Budzyńska et al., 2017; Igarashi et al., 2014). We next analyzed expression profiles of cytokine receptors, signaling by which is known to influence CSR. External signaling from IL-4 is known to drive IgG1 switching (Higgins et al., 2019). We observed up-regulation of the cognate receptor, *Il4ra* (Figure 4F), which suggests a model where increased sensitivity to IL-4 signaling may sustain commitment to IgG1 switching. In contrast, transcripts for the cytokine receptors *Ifnγr1* and *Il5ra* were not substantially changed in the inferred trajectory (Figure 4G,H). IFN-γ signaling has been demonstrated to inhibit IgG1 switching (Kawano et al., 1994) whereas the role of IL-5 in CSR has remained controversial (Huston et al., 1996; Matsumoto et al., 1989; Purkerson and Isakson, 1992). In conclusion, Pseudocell tracing revealed increased expression of *Il4ra, Bach2 and Aicda*, importantly after CSR has been initiated. *Il4ra* may function to reinforce IL4 signaling which directs CSR to the IgG1 isotype, whereas *Bach2* may upregulate *Aicda* expression thereby enabling efficient SHM after CSR.

## DISCUSSION

We present a supervised machine learning framework, Pseudocell Tracer, for modeling cellular trajectories in complex systems. Inference of cellular trajectories from scRNA-seq datasets remains a challenging problem, particularly for heterogeneous cellular compartments with complex dynamics. Existing methods for trajectory inference are strongly dependent on initial low-dimensional projections of the datasets. Concurrent and inter-digitated transcriptional programs, such as those regulating cell cycle and metabolism can obstruct unsupervised dimensionality reduction and pseudotemporal ordering techniques. This problem can be amplified by sparse sampling of key intermediate cellular states. To address these two challenges, we harness adjacent biological information in order to shape the latent space in a biologically valid manner thereby revealing discrete cellular trajectories in a complex developmental compartment that are otherwise obscured.

In this work we also introduce and propose a new paradigm for characterizing cellular trajectories through pseudo cell state space rather than time. In particular, Pseudocell Tracer is able to generate pesudocells at regular intervals along a virtual cell-state axis. By harnessing recent advances in deep generative modeling, Pseudocell Tracer learns a generative model encompassing all cells in a biologically meaningful latent space. As a result, the generative model provides a means to interpolate cells in the latent space and allows for the specific delineation of pseudocells by conditioning on adjacent biological information. Importantly, we demonstrate that even a relatively small dataset with a few hundred cells is sufficient for learning and can be used to generate biologically plausible virtual cells. The surprising effectiveness of GANs in simulating realistic subpopulations of cells from small datasets has been independently demonstrated (Marouf et al., 2020).

Pseudocell Tracer was used to analyze and infer gene expression dynamics along a particular CSR trajectory (IgG1), during a prototypic antigen-induced B cell response. In spite of extensive genetic and molecular analysis of CSR, the gene expression dynamics of B cells undergoing CSR in vivo have not been revealed. In fact, a recent report using extensive scRNA-Seq profiling of human tonsillar B cells was still unable to reveal the developmental modulation of genomic states underlying particular CSR trajectories using unsupervised dimensionality reduction techniques (King et al., 2020). We utilized a unique scRNA-seq dataset generated from antigen-specific B cells induced by NP-KLH immunization to infer the cellular trajectories of germinal center B cells undergoing CSR to IgG1, the dominant isotype manifested under these conditions. Although recent work has indicated that CSR is primarily induced before entry of antigen-specific B cells into germinal centers (Roco et al., 2019), we were able to detect it also within the GC compartment.

Our results revealed an ordering of key transcription factors regulating CSR, including the concomitant induction of *Nfkb1* and *Stat6* prior to the upregulation of *Bach2* expression. The former transcription factors have known roles in regulating IgG1 CSR and the latter regulates *Aicda* gene expression and therefore both CSR and SHM. An intriguing finding is the upregulation of *Aicda* gene expression in the CSR trajectory after it has been initiated suggests that increased expression of AID maybe needed for efficient SHM which occurs in the germinal centers. Finally, the expression dynamics of genes encoding cytokine receptors point to the existence of a regulatory mechanism that reinforces IL-4 signaling to direct CSR to the IgG1 isotype. Although we demonstrate proof-of-concept in modeling CSR along a particular isotype trajectory, we anticipate future studies with larger high-throughput scRNA-seq datasets will capture more cells, particularly representing other isotypes and facilitate assembly of isotype-specific B cell trajectories in diverse lymphoid organs and tissues. In the current work, we utilized an encoding that reflected potential CSR paths, however, Pseudocell Tracer can encode other structured adjacent biological information as well, such as phylogenetic trees constituted by somatically mutating antibody variable regions. In so doing, Pseudocell Tracer could be used to guide the latent space and conditional generation of specific trajectories of B cells undergoing SHM and affinity maturation.

The machine learning framework underlying Pseudocell Tracer provides for a flexible means to explore new forms of adjacent biological information in extremely diverse contexts. Ultimately, Pseudocell Tracer is a powerful framework for characterizing the transcriptional states and trajectories of cells during their development and activation. These states and trajectories, particularly rare ones, can be revealed by embedding them in valid biological priors. For example, Pseudocell Tracer has the potential to merge multiple single cell profiling experiments from different biological compartments, providing a novel way to bridge datasets by using prior molecular information about their relatedness. Therefore, Pseudocell Tracer promises to be a robust engine for hypothesis generation for experimental biology by predicting novel regulators and rare cell states underlying extremely diverse cellular trajectories.

## MATERIALS AND METHODS

### Mice and immunization

C56BL/6J (Jax 000664) mice were obtained from the Jackson Laboratory. Mice were housed in specific pathogen-free conditions and were used and maintained in accordance of CCHMC Institutional Animal Care and Use Committee guidelines. Six to eight week old mice were immunized intraperitoneally with 100 µg NP(23)-KLH (Biosearch Technologies) mixed with 50% (v/v) Alum (Thermo Scientific) and 1 µg LPS (Sigma).

### Flow cytometry and B cell sorting

Splenocytes were washed and prepared as single-cell suspensions in MACS buffer (pH 7.4; PBS plus 1% FBS and 5 mM EDTA). Nonspecific antibody binding was blocked with 2.4G2 (BD, 25 µg/ml) by incubation for 15 min on ice. Cells were stained for 30 min at 4 °C with indicated antibodies. All antibodies and relevant reagents used for flow cytometry are listed in Extended Data Table 1. Data were collected on LSRII or Fortessa 2 (BD) and were analyzed with FlowJo software (TreeStar).

Single cell suspensions were prepared from mouse spleens in MACS buffer on day 13 post immunization and blocked with 25 µg/ml 2.4G2 (BD) for 15 min on ice. Cells were then labeled with 2 µg/ml biotin anti-CD3, 1 µg/ml biotin anti-CD4, 1 µg/ml biotin anti-CD8, 2 µg/ml biotin anti-CD11C and 2 µg/ml biotin anti-IgD (Extended Data Table 1) for 20 min at 4°C. After washing three times, cells were incubated with anti-biotin beads (Miltenyi Biotec) for 20 min at 4°C. Magnetic columns were used to deplete non-B cells according to standard protocol (Miltenyi Biotec). Cells in effluent were further labeled with NP-PE, B220, Fas and GL-7 antibodies (Extended Data Table 1) for 30 min at 4 °C. GC B cells were sorted as 7AAD–B220+FashiGL-7hiNP+ using FACSAria II (BD) with 70 µm nozzle at 4 °C.

### scRNA-Seq Data Generation

Single B cells were prepared using the C1TM Single-Cell Auto Prep System (Fluidigm) according to the manufacturer’s instructions. In short, flow-sorted cells were re-suspended at a concentration of 3×105 cells/ml then loaded onto a primed C1 Fluidic Chip for mRNA-Seq (5-10 µm). Cell separation was visually scored, 44-61 single cells were captured in each run. Cells were lysed on chip and reverse transcription was performed using Clontech SMARTer® Kit using the mRNA Seq: RT + Amp (1771x) according to the manufacturer’s instructions. After reverse transcription, cDNAs were transferred to a 96 well plate and diluted with C1TM DNA Dilution Reagent. Quant-iTTM PicoGreen® dsDNA Assay Kit (Life Technologies) and Agilent High Sensitivity DNA Kit (Agilent Technologies) were used to quantify cDNAs. Libraries were prepared using Nextera XT DNA Library Preparation Kit (Illumina) using cDNAs with an initial concentration>200 pg/µl, diluted to 100 pg/µl. In each single-cell library preparation, a total of 125 pg cDNA was tagmented at 55 °C for 20 minutes. Libraries were pooled and purified on AMPure® bead-based magnetic separation before a final quality control using Qubit® dsDNA HS Assay Kit (Life Tecnologies) and Agilent High Sensitivity DNA Kit. 96 scRNA Seq libraries were sequenced per lane on HiSeq 2500 with 75bp paired-end sequencing (∼300 million bp/gel).

### scRNA-Seq Data Preprocessing

Genes containing no counts across any samples were discarded. We calculated the relative isotype expression within a B cell by dividing the log2(TPM + 1) expression of each distinct isotype transcript with the cumulative expression of all isotypes. The relative isotype expression profiles were clustered using hierarchical clustering using the *clustergram* function in Matlab with default settings. For training the autoencoder and CGAN models, the count data of the four batches were normalized using the *scran* R package’s *MultiBatchNorm* function, resulting in rescaled log-counts for each gene (Lun et al., 2016). Further batch correction was then applied using the Mutual Nearest Neighbor (MNN) approach from the *batchelor* R package (Haghverdi et al., 2018), resulting in MNN corrected relative expression values.

### Pseudotime Trajectory

We apply three state-of-the-art pseudotime trajectory inference methods to our data: Monocole3, PAGA, and Slingshot. Monocle3 is a method to learn pseudotime through the use of dimension reduction and graph learning. A minimum spanning tree is constructed and used to order the cells to infer pseudotime trajectory. We implemented Monocle3 using the default settings with UMAP dimension reduction and Louvain clustering. PAGA works by constructing a k-nearest neighbor graph, which is then clustered into modules using the Louvain algorithm and connectivity is assigned between the modules. We implemented PAGA using default settings and removed connections less than 0.1. Slingshot is another single cell trajectory inference method. In their method, they project the data into a latent space and perform clustering. In our implementation of Slingshot, we use PCA for dimension reduction and Gaussian mixture model (GMM) clustering with default parameters. The minimum spanning tree is then constructed from the clusters.

### Supervised Autoencoder

In order to shape the latent space in a meaningful way, we utilize a supervised autoencoder to help distinguish important differences in cell subtypes. Our work constructs a supervised autoencoder in two steps. The first step employs supervised encoding using the relative abundance of the IgH genes of interest based on their relative expression. To do so, the normalized gene counts are encoded into a latent layer using a neural network model, and this latent layer is then used to predict the relative abundance of the different IgH genes. The encoder has an input of size 41,671 and contains two fully connected layers between the input and the latent layer of sizes 512 and 256 respectively, each using the rectified linear unit (ReLU) activation function. The latent layer has a size of 64 nodes and uses the sigmoid activation function. In addition, there is a fully connected layer of size 128 between the latent layer and the output. The output layer contains 8 nodes and uses the softmax activation function in order to generate a relative distribution across the 8 IgH genes. The network is trained using the Adam optimizer with a learning rate of 1×10^−4^ and the Kullback-Leibler divergence (KLD) loss function,

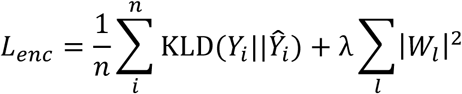

Here, the first term represents the average KLD between the observed IgH proportions, Y_*i*_, and the predicted IgH proportions, Ŷ_*i*_, across all *n* samples. The second term regularizes the network using the weighted sum the L2-norms for the weights of each layer *W* _*l*_, *where l* represents the layer. In our study, we found the best weight coefficient to be λ = 1×10^−5^. Further regularization is added by implementing dropout with a rate of 0.3 at every hidden layer. The network is trained using early stopping, where 10% of the training data is held out as a validation set. The KLD term of the loss function was evaluated on the validation set each epoch and training was terminated if there had not been a decrease in 100 epochs, whereafter the model was reverted to the previously best state. The model is then trained for an additional 5 epochs using the entire training data, including the validation set that was used for early stopping.

After the supervised encoder had been trained, the second step employed a decoder in order to reconstruct the original normalized gene values from the latent space. This network takes the latent space of size 64 as an input to predict the normalized gene expression for each of gene as an output. The network contains two fully connected layers of sizes 256 and 512 respectively, each using the ReLU activation function. The output layer has a size of 41,671 and uses the linear activation function. The network is trained using the Adam optimizer with a learning rate of 1×10^−4^ with the mean squared error (MSE) loss function,

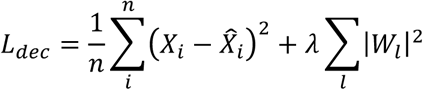

Here, the first term represents the average MSE between the observed normalized gene expression, *X*_*i*_, used to generate the latent space of some sample *i*, and the decoded normalized gene expression 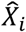. The second term regularizes the network using the weighted sum the L2-norms in the same way as seen in the supervised encoder. We again found the best regularization parameter to be λ = 1×10^−5^. The decoder model was further regularized using batch normalization at each of the hidden layers. The model was trained using the same early stopping approach from the supervised encoder, however in this case we stop the training based on the MSE term of the loss function.

### Conditional Generative Adversarial Network

In order to generate samples for unobserved pseudo-times, we utilize a CGAN architecture (Goodfellow et al., 2014). A CGAN is composed of two networks: a generator and a discriminator. The generator model learns to generate fake data that is as close to the distribution of the real data as possible. At the same time, the discriminator model tries to predict if a piece of data is real or fake. Both models are trained in an adversarial manner where the generator tries to maximize the log-probability of labeling real and fake images correctly while the discriminator tries to minimize it, resulting in a zero-sum minimax game. The models are trained using gradient descent until Nash equilibrium is reached.

The generator in our model, *G*, takes in as an input a vector of 32 values sampled from ∼U(−1,1) concatenated with the relative abundance of the 8 IgH genes in order to output a vector representing the encoded latent representation of gene expression values. The network contains two fully connected layers of sizes of 256 and 512 respectively, both using the ReLU activation function. The output is size 64 with a linear activation function. The discriminator, *D*, takes a vector of size 64 representing encoded gene expression as well as the relative abundance of the 8 IgH genes and outputs a single value between 0 and 1 as the probability of the data being real. The discriminator has a single layer of size 512 that uses the ReLU activation function. To avoid problems of a vanishing gradient, we adjust the loss function of the generator to create a non-saturating loss function (Goodfellow et al., 2014). The networks are trained simultaneously using the Adam optimizer with a learning rate of 5×10^−5^ until convergence (about 50,000 epochs). The loss function for the discriminator and generator are shown below.

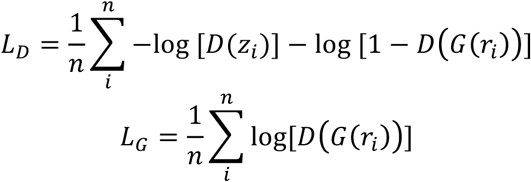

Here *z*_*i*_ represents the latent space generated from the observed gene expression from sample *i* using the and *G*(*r*_*i*_) represents the generated latent space using the relative IgH values from sample *i*. Latent spaces are taken from the autoencoder model trained on the entire data set.

### Pseudocell Generation

Once the CGAN model was trained to equilibrium, we generated 100 latent spaces for each relative isotype expression profile. For each of IgG1, IgG2b, and IgG3, we generated latent spaces for each trajectory starting at 100% IgM, and for each subsequent relative isotype expression profile, we reduced the relative abundance of IgM by 1% and increased the respective isotype gene expression by 1%. We then decoded each latent space back to the normalized gene expression values in order to obtain the gene trajectories. For visualization, the 95% confidence interval is plotted along with averages. The relative expression of each gene shares the same space as the NMM corrected log-scale gene expression.

### Data and Code Availability

All raw single cell RNA-seq data from this work is submitted to the GEO repository: GSEXXXXX. Software code used in generating the results is described above in detail and on GitHub: https://github.com/akds/pseudocell

## SUPPLEMENTRY FIGURE LEGENDS

Supplementary Figure 1: t-SNE of RNA-seq data, colored by cell cycle genes.

Supplementary Figure 2: Detailed architecture of the supervised autoencoder.

Supplementary Figure 3: Detailed architecture of the conditional GAN.

Supplementary Figure 4: Plot of relative expression of *Aicda, Ighm* and *Ighg2b (left) and Ighg3 (right)*.

**Extended Data Table 1.** Reagents used for flow cytometry.

| **Antibodies** |  |  |  |
| --- | --- | --- | --- |
| **Target** | **Clone** | **Label** | **Provider** |
| B220 | RA3-6B2 | Brilliant Violet 510, Alexa Fluor 700, FITC | Biolegend |
| GL-7 | GL-7 | FITC | Biolegend |
| IgG1 | RMG1-1 | PE-CY7 | Biolegend |
| IgD | 11-26c (11-26) | eFlour450, Biotin | eBioscience |
| CD11C | N418 | Biotin | eBioscience |
| CD4 | GK1.5 | Biotin | eBioscience |
| CD8 | 53-6.7 | Biotin | eBioscience |
| CD3 | 145-2C11 | Biotin | eBioscience |
| IgM | eB121-15F9 | eFlour450, eFlour660 | eBioscience |
| Fas | Jo2 | PE, PE-CY7 | BD Bioscience |
| GL-7 | GL-7 | Brilliant Violet 421 | BD Bioscience |
| **Dyes and reagents** |  |  |  |
| Fixable Viability Dye eFluor 780 |  |  | eBioscience |
| Fixable Viability Dye eFluor 450 |  |  | eBioscience |
| 7AAD |  |  | BD Bioscience |
| NP(31)-PE |  |  | Biosearch Technologies |

